# Positively interacting strains that circulate in a network structured population induce cycling epidemics of Mycoplasma Pneumoniae

**DOI:** 10.1101/312066

**Authors:** Xu-Sheng Zhang, Hongxin Zhao, Emilia Vynnycky, Vicki Chalker

## Abstract

In many countries *Mycoplasma pneumoniae* (MP) epidemics last approximately one to two years and occur every three to seven years. Poor understanding of the drivers of recurrent MP epidemics limits the predictability of and dynamic responses to the outbreak. Taking into account network structured contacts among people and co-circulating strains of MP, we propose a multi-strain SIRS network model of epidemics of MP where different strains interact during re-infection and within secondary infection. Simulations show that although strain interactions and network-mediated spatial correlations are two separate mechanisms for MP epidemics cycling, each requires very restricted model parameter values such as strong strain interactions and strong network contacts, respectively. When both mechanisms work collectively, MP recurrent epidemics become feasible within the plausible ranges of model parameters. This indicates that positively interacting strains that co-circulate within network contacts induce periodicity and dominant strain shift in observed MP incidence.

## Introduction

*Mycoplasma pneumoniae* (MP) is an “atypical” bacterium that causes acute respiratory infection in humans of all ages. *M. pneumoniae* is considered a common cause of pneumonia: MP causes about 15–20% adult community-acquired pneumonia (CAP) and up to 40% cases in children; however, not every infected patient actually develops pneumonia (Foy, 1993; Korppi *et al*., 2004; Dumke *et al*., 2012). MP infection generally tends to occur more frequently during the summer and autumn months when other respiratory pathogens are less prevalent; but the disease incidence does not appear to be related to season or geography (Waites and Talkington 2004; Winchell, 2013). For example, we also notice that MP infections have been observed to occur more frequently in winter months in England and Wales (Chalker *et al*., 2011; Brown *et al*., 2016). Epidemics of MP tend to occur every 3-7 years in the general population (Chalker *et al*., 2011; Jacobs, 2012; Brown *et al*., 2016). Analysis of laboratory reports of MP infections in England and Wales from 1975 to 2009 (Nguipdop-Djomo *et al*., 2013) has indicated that these epidemics last on average 18 months occurring at approximately four yearly intervals. *M. pneumoniae* is a polymorphic pathogen (Dorigo-Zetsma *et al*., 2000; Pereyre *et al*., 2012): for example, Chalker *et al*. (2011) identified eleven strain types circulating in England and Wales during October 2010 to January 2011. MP strains can be differentiated based on differences in the P1 adhesin gene or in the number of repetitive sequences at a given genomic locus using multilocus variable number tandem repeat analysis (MLVA) (Dumke and Jacobs, 2011; Simmons *et al*., 2013). Kenri et al. (2008) noticed that more than one serotype of MP were circulating within Japanese populations. Kogoj *et al*. (2017) observed a shift in the dominant MP strain between two epidemics that occurred in Slovenia in 2006 and 2016. Multiple strains of MP and their co-circulation were also observed in other countries (e.g., Dumke *et al*., 2010; Spuesens *et al*., 2009; Martinez *et al*., 2010; Zhao *et al*., 2015; Brown *et al*., 2016). Although there are many different isolates and strains, analysis of repetitive elements distributed in variable size and sequence over the genome of MP strains suggested two main types: P1 type 1 and P1 type 2 (Kenri *et al*., 2008; Spuesens *et al*., 2009; Brown *et al*., 2015; Dumke and Jacobs, 2016).

Humans are the sole reservoir of MP and transmission requires close contact. Outbreaks typically occur within closed populations, such as in schools, military premises and prisons. Airborne spread of aerosols and, potentially, indirect contact with contaminated items, may contribute to transmission. The transmissibility of an infectious agent can be estimated by calculating the basic reproduction number (R0), which is defined as the mean number of secondary infectious cases generated by one primary infectious case introduced into a totally susceptible population (Anderson and May, 1991). Using seroprevalence data from a western population, Nguipdop-Djomo *et al*. (2013) estimated *R*_0_ of MP to be 1.7 (95% CI 1.6–1.9), indicating low transmissibility. The incubation period of MP averages 2 to 3 weeks. The duration of infectiousness is unclear and is commonly estimated to be up to 3 weeks from onset of illness (Clyde, 1993). Immunity occurs post infection, but later re-infection with different subtypes is recognized, suggesting the immunity is not lifelong and no strong cross protection between different subtypes (Foy *et al*., 1977; Ito *et al*., 2001; Dumke and Jacobs, 2016). The duration of immunity ranges from 2 to 10 years (Lind *et al*., 1997; Omori *et al*., 2015).

Seasonal forcing in transmission has been proposed as one determinant for the periodic patterns in other infectious diseases (Keeling and Rohani, 2008); however, Omori *et al*. (2015) found that the seasonal forcing that occurs annually cannot generate the multi-year periodicity of MP incidence. They (Nakata and Omori, 2015; Omori *et al*., 2015) further proposed that the certain finite delay in the progression from immunity to the susceptible may provide an explanation to the occurrence of the cyclic epidemics of MP infections. More concretely, Omori *et al*. (2015) show that “minor variation in the duration of immunity at the population level must be considered essential for the MP epidemic cycle because the MP cyclic incidence pattern did not replicate without it.” As shown in Figure 3 of Omori *et al*. (2015), this requires that the distribution for the duration of immunity should have a variance of around 0.63. Up to now no empirical data are available for estimating the distribution of the duration of MP immunity.

**Figure 3.**
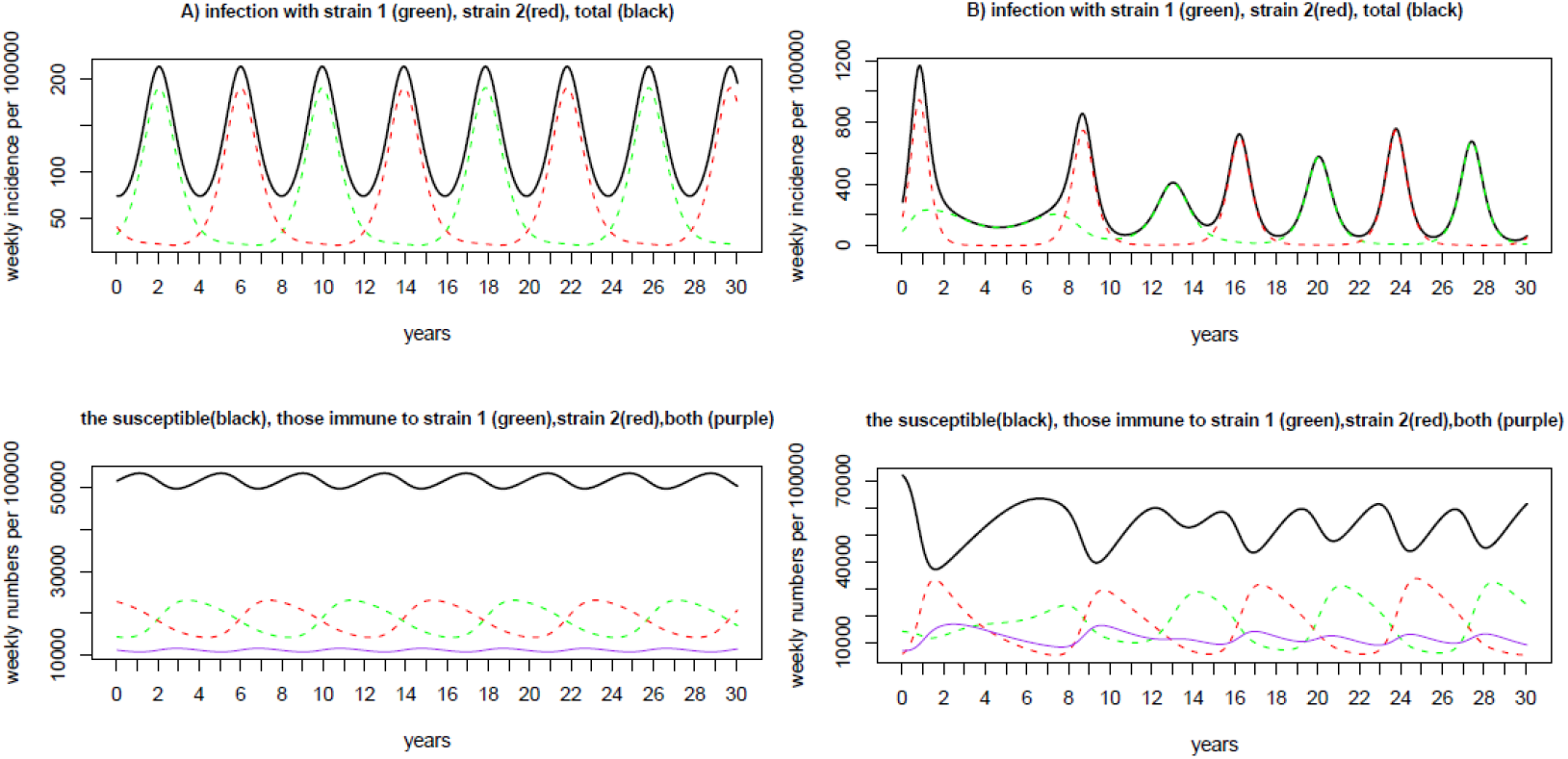
Two examples of epidemic curve from LHS samples under the situation of no interaction within the secondary infection (i.e., *ν*=*μ*=1). Panel A) *κ*=5.69, *ψ*=0.415, *D* = 17.2 days and *d* = 9.2 years, EpiT =4 years, CV =0.51; panel B) *κ*=3.34, *ψ*=0.202, *D* =37.7 days, *d* = 4.1 years, and EpiT =4 years, CV =0.93. The legend is provided in the title to each figure. It is obvious that the proportion of individuals that are simultaneously immune to both strains is kept low in the two examples. The proportion of individuals immune to one strain alone is temporally highly anti-correlated with the proportion of these immune to the other strain alone, with the correlation coefficient −95% and −84% for panel A) and panel B) respectively.

The MP incidence in England and Wales has declined (Brown *et al*., 2016) following the widespread use of macrolides antibiotics since introduction in the late 1990s’ (Woodhead and Macfarlane, 2000). Due to the emergence of macrolide-resistant strains, MP infections are of increasing public health interest (Morozumi *et al*., 2008; Zhao *et al*., 2013). An understanding of the mechanisms by which recurrent epidemics of MP infection occur is urgently needed to enable control of future epidemics.

The two distinctive aspects of the MP epidemics: the prevalent serotype shifts among epidemics (Kenri *etal*., 2008; Suzuki *et al*., 2015; Zhao *et al*., 2015; Brown *et al*., 2016; Kogoj *et al*., 2017) and cycling of MP incidence may be interconnected. This has been proposed before. Dumke *et al*. (2010) and Spuesens *et al*. (2009) argued that MP epidemics arise due to a change in the two main P1 types and variants of P1 sequences. Chalker et al (2011) observed increased incidence of MP infection correlating with co-circulation of multiple strains within the population of England and Wales. Brown *et al*. (2016) speculated that dominant strain shift may be the cause of recurrent MP epidemics in view of the presence of multiple strains in observed increases of MP infection. Despite a lack of current data (due to limited focus on MP internationally and poor tools for detection and simultaneous strain discrimination) we speculate that serotype interactions such as synergistic associations and competition, in addition to the cross-immunity of differing P1 types, exist and play a possible role in the recurrent epidemics of MP infections. Previous transmission dynamics models (see the review of Omori *et al*. 2015) neglected the following phenomena: co-circulation within human populations of multiple strains of MP and network structural contact patterns among people. Infection transmission depends on the contact rate as well as whom each individual contacts. Recent studies (Mossong *et al*., 2008) showed that people do not mix randomly. For example, contact patterns between people may display the characteristics of scale-free networks (Pastor-Satorras and Vespignani, 2001) or small-world networks (Watts and Strogatz, 1998). An important parameter of a network is its degree, defined as the number of other individuals to which one is connected. A well-mixed network (i.e., the loose network) will have a high average degree while a less mixed network should have a small average. Realistic networks of contacts that are relevant to infectious diseases usually have a small average degree (Leventhal *et al*., 2015). On the contrary, the assumption of random mixing, in which every person is equally likely to contact any other person within the population (Keeling and Rohani, 2008; Diekmann *et al*., 2013), results in a very large degree.

Network structured models describe the transmission dynamics as in spatial transmission processes among connected groups and thus induce spatial correlation between infections. Letting infection spread on a homogeneous population with a fixed random network structure, Rozhnova and Nunes (2009) illustrate that this spatial correlation within Susceptible-Infectious-Recovered-Susceptible models assists the generation of sustained cyclical epidemics. However, strong spatial correlation (i.e., strong network structure) was needed for the cycles to persist when they just considered the transmission dynamics of a single strain in a population. Considering a two strain version of the SIRS epidemic network model (Zhang, 2016), the restriction on model parameters especially the degree of contacts is much relaxed. Recurrent epidemics were also predicted by models in a population which did not have a network structure, but in which people could be re-infected or co-infected with multiple strains (Zhang and Cao, 2014). Neither of these studies considered MP infection and we explore whether inclusion of both factors – a) competition between strains in a network-structured population and b) re-infection and co-infection with multiple strains - can explain the observed cycles in MP incidence

## Models and Methods

### General structure of the model

We consider a Susceptible-Infectious-Recovered-Susceptible, rather than a Susceptible-Exposure-Infectious-Recovered-Susceptible structure that has been used in other studies (Omori *et al*., 2015). This simplicity is justified as we focus on the long term behaviour of MP transmission dynamics, and the exposure stage does not influence the overall transmissibility and long-term patterns (Diekmann *et al*., 2013; Omori *et al*., 2015). Since the many different isolates and strains of MP can be classified into two main types: P1 type 1 and P1 type 2 (Kenri *et al*., 2008; Spuesens *et al*., 2009; Brown *et al*., 2015; Dumke and Jacobs, 2016), our model just considers the transmission dynamics of two strains. Within the SIRS transmission dynamics model, a population of size *N* is modelled as a network in which every individual randomly contacts a fixed number (*κ*) of other individuals, and is classified into eight compartments (Figure 1), namely those who are susceptible to infection with any strain (S), those who are infected and infectious with strain 1 or 2 (I_1_ and I_2_), those who have recovered from infection with a given strain and are susceptible to infection with the other strain (R_1_ and R_2_), those who are infected and infectious with strain 1 or 2 after recovering from previous infection (J_1_ and J_2_) and those who are immune to infection with both strains (R). We refer to people in the I_1_ and I_2_ compartments as those with “primary infection”, and to people in the J_1_ and J_2_ compartments as those with “secondary infection”. Individuals are denoted by nodes and contacts between individuals by edges.

**Figure 1.**
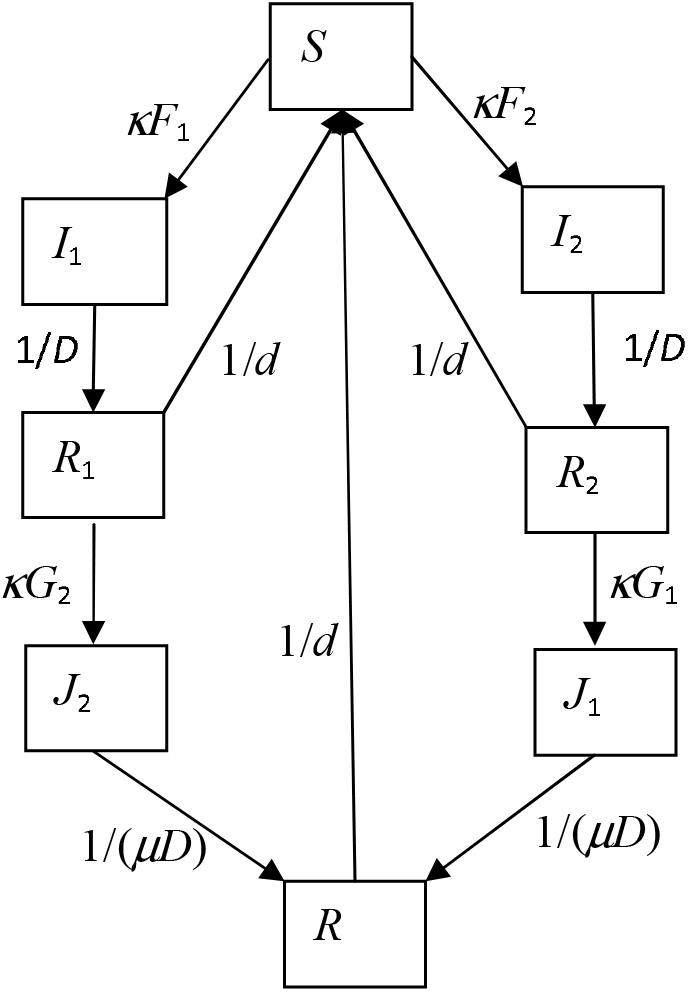
Flow chart of the two-strain SIRS epidemic model. Arrows indicate transitions. Expressions next to arrows show the *per capita* flow rate between compartments. Births and deaths are not shown. Parameter *κ* is the degree of contacts each person has and *μ* is the effect of primary infection strain on the duration of infection by the secondary strain. Variables *F*_1_ (*F*_2_) and *G*_1_ (*G*_2_) are forces of infection of strain 1 (strain 2) that are defined in equations (4).

The epidemic dynamics is determined by the following transmission and transition processes. Susceptible nodes (*S*) become infected with strain *i, i* = {1,2}, at rate *λ* through an edge with a node of primary infection *I_i_*., or at rate *νλ* through an edge with a node of secondary infection *J_i_*. Here parameter *λ* represents the constant transmission rate and parameter *ν* is the relative infectiousness of a secondary infection, compared to a primary. Primarily infected nodes (*I_i_*) stay infectious on average for *D* days before becoming fully immune (*R_i_*) to the infecting strain *i* and partially so to the other strain. Recovered individuals (*R_i_*) stay immune for an average of *d* days before becoming susceptible again, or becoming secondarily infected at rate (1-*ψ*)*λ* through an edge linked with a node of infection (*I*_3-*i*_ or *J*_3-*i*_) to become secondarily infected *J*_3-*i*_, *i*={1,2}. Here *ψ* reflects the reduction in susceptibility due to previous exposure to the other strain (i.e., cross-immunity). Nodes of secondary infection *J_i, i_* = {1,2} stay infectious for an average of *μD* days before becoming fully immune to all strains (i.e., *R*). Here parameter *μ* defines the effect of having experienced primary infection on the duration of the secondary infection. Nodes of *R* stay fully immune for an average of *d* days before becoming susceptible again. Therefore naïve individuals are recruited into the population through birth and loss of immunity. These transitions and transmissions are defined according to the pairs or triplets involved in the process (Eames and Keeling, 2002; Rozhnova and Nunes, 2009). For simplicity we ignore clustering in the network (c.f., Eames and Keeling, 2002; Leventhal *et al*., 2015).

### Model equations

Similar to Eames and Keeling (2002), the proportions of people in eight compartments are represented by [S], [I_1_], [I_2_], [R_1_], [R_2_], [J_1_], [J_2_], and [R]. Because of the constant population size (i.e., the constant number of nodes), [S] +[I_1_] +[I_2_] +[R_1_] +[R_2_] +[J_1_] +[J_2_] +[R] =1. There are (8×7)/2 = 28 heterogeneous pairs within the network in which the two nodes of a pair are of different states. The proportion of the population that is in a pair ([XY]) is defined as

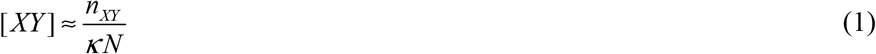

Here *n_XY_* is the number of pairs within the population. The number of homogenous pairs can be found from these equations for heterogeneous pairs: e.g., 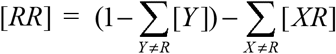 and 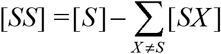. The state of the model system is defined by eight integers of nodes and 28 integers of heterogeneous pairs. To focus on the impact of spatial correlation mediated by network structure (i.e., competition among the limited number of partners) and interactions between strains, two strains are simply assumed to be antigenically indistinguishable within linked patients.

Transmission of infection among nodes occurs through pair-link and the change of pairs is determined by the triples. To close the model system, the proportion, [*XYZ*], of the triple *XYZ* with node *Y* having contacts with both *X* and *Z* is approximated in terms of the proportion of pairs as in Eames and Keeling (2002),

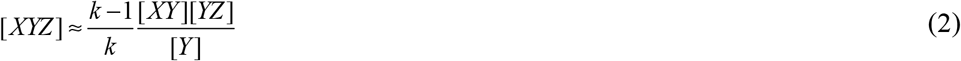

The flow chart of the transmission dynamics model is shown in Figure 1. The standard pair approximation SIRS model of two strains is described by a set of 28 + 7 = 35 differential equations as,

**Equations describing the time changes in 7 nodes**

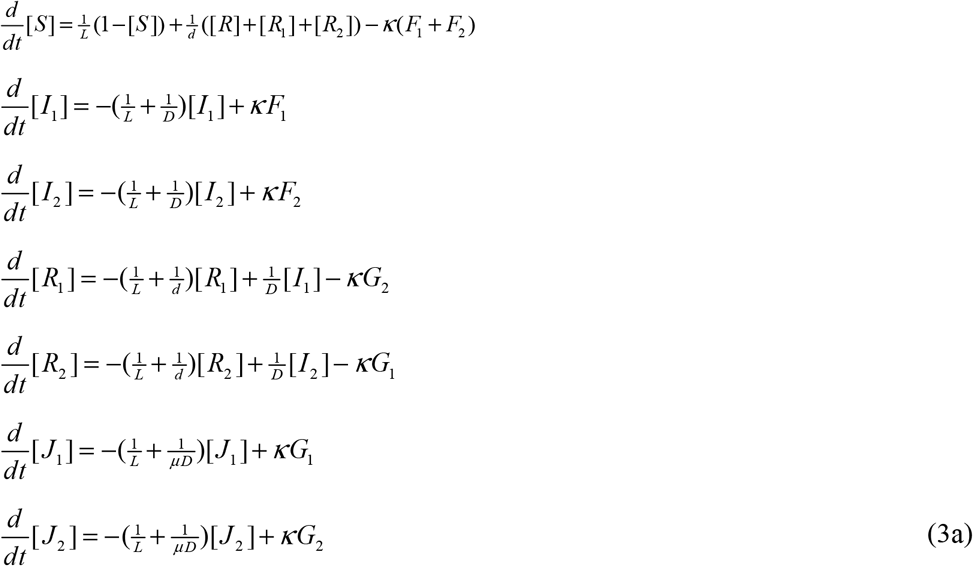

**Equations describing the time changes of 28 pairs**

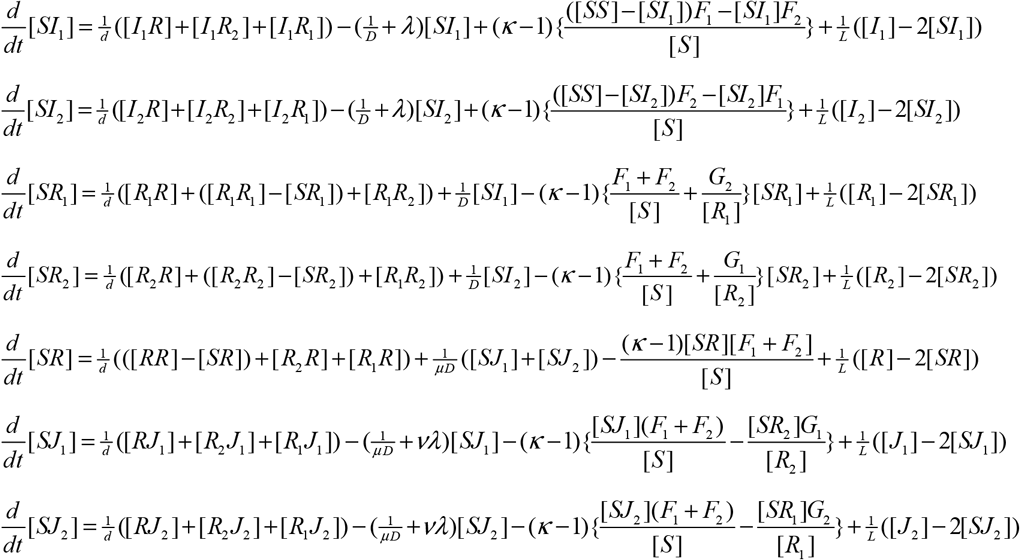

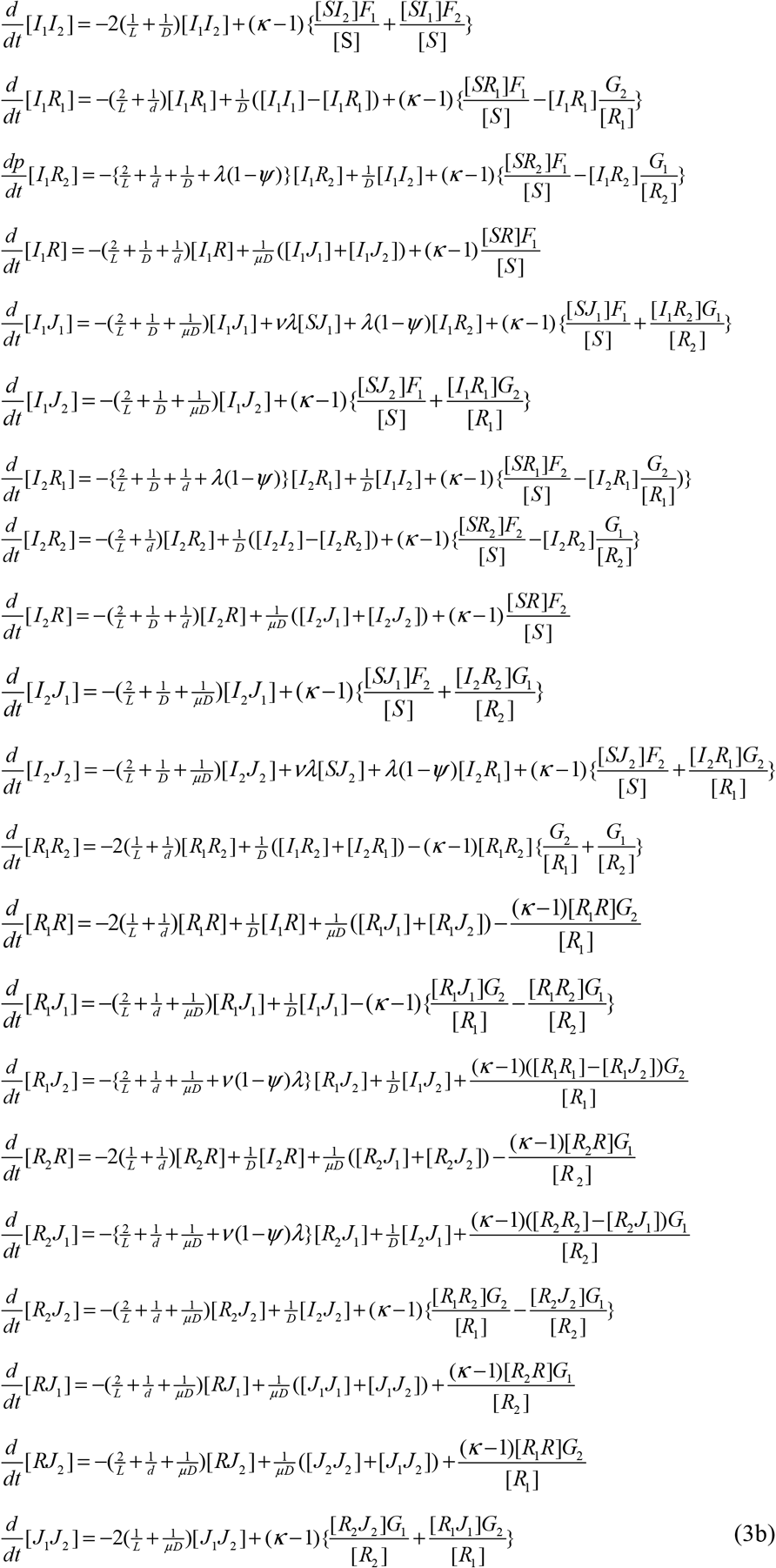

In the above equations, the different forces of infection are

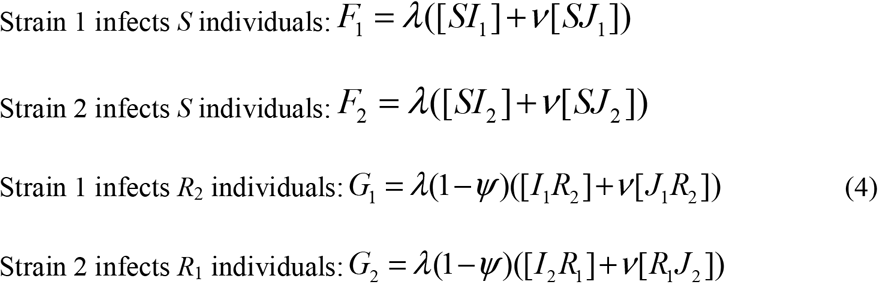

The parameters of the model system are defined in Table 1. Compared with the model presented in Zhang (2016), here we introduce two interaction parameters to define the effects of experiencing primary infection on infectivity (*ν*) and the duration (*μ*) of a secondary infection. The complexity of the two strain network dynamics allows us to investigate the combined effects of strain interactions (cross-immunity during re-infection and effects of the primary infection on a secondary infection) on dynamic patterns of endemic infectious diseases, along with spatial competition embedded within the random network. Ignoring the stochasticity due to the limited size of population, here we focus on these by considering an infinitely large population (i.e., *N*→∞).

**Table 1.** Model parameters.

| parameter | Definition | Values or ranges | source |
| --- | --- | --- | --- |
| $L$ | Average life span | 70*365 days | The World Fact-book Life Expectancy |
| $D$ | Infectious period of a single infection | 14 – 42 days | Clyde, 1993;Omori *et al.*, 2015 |
| $d$ | Duration of immunity | 2 – 10 years | Lind *et al.*, 1997 |
| $R_0$ | Basic reproduction number | 1.7 | Nguipdop-Djomo *et al.*, 2013 |
| $\lambda$ | Transmission rate of single infection | $\lambda = \frac{R_0(d+1)}{D[d(k-2) + (\kappa-1)]}$ | Eames and Keeling, 2002 |
| $\psi$ | Reduction in susceptibility to a secondary infection, resulting from a primary infection (“cross-immunity”) | 0.0 – 1.0 | – |
| $\nu$ | Relative infectiousness of a secondary infection, compared to a primary infection. | 0.5 – 2.0 | Negative (<1) and positive (>1) effects |
| $\mu$ | Factor by which the duration of a secondary infection differs from that of a primary infection. | 0.5 – 2.0 | Negative (<1) and positive (>1) effects |
| $\kappa$ | Degree of contact network: number of people with whom one person has contact. | 2.5 – 25.0 | assumed |

## Methods

We used Latin hypercube sampling (Iman *et al*., 1981) to identify parameter values which led to model predictions of cycles in incidence which were consistent with those observed. The values of the following parameters were sampled within the ranges listed in Table 1: infectious period (*D*), duration of immunity (*d*), degree of contact network (*κ*), cross-immunity (*ψ*), effects of primary infection on infectivity (*ν*) and duration (*μ*) of a secondary infection. This was done with the function randomLHS of the package lhs in R computing language (R Development Core Team, 2015).

The basic reproduction number (*R*_0_) was fixed at 1.7 as estimated by Nguipdop-Djomo *et al*. (2013). When estimating *R*_0_, Nguipdop-Djomo *et al*. (2013) assumed random mixing among individuals and didn’t account for a network structured population. For a network structured population, the random mixing assumption will give rise to an over-estimate of *R*_0_ (Figure 2 of Eames and Keeling, 2002), so we also consider this effect by assuming that *R*_0_= 1.5 and 1.3 for MP. The total number of infections in our model includes both asymptomatic and symptomatic infections. Since MP may affect all age groups (e.g., Ito *et al*., 2001; Chalker *et al*., 2011; Brown *et al*., 2016), we consider the situation in which the life span is 70 years, the worldwide average life expectancy according to the world fact book (<u>The World Fact-book Life Expectancy</u>).

**Figure 2.**
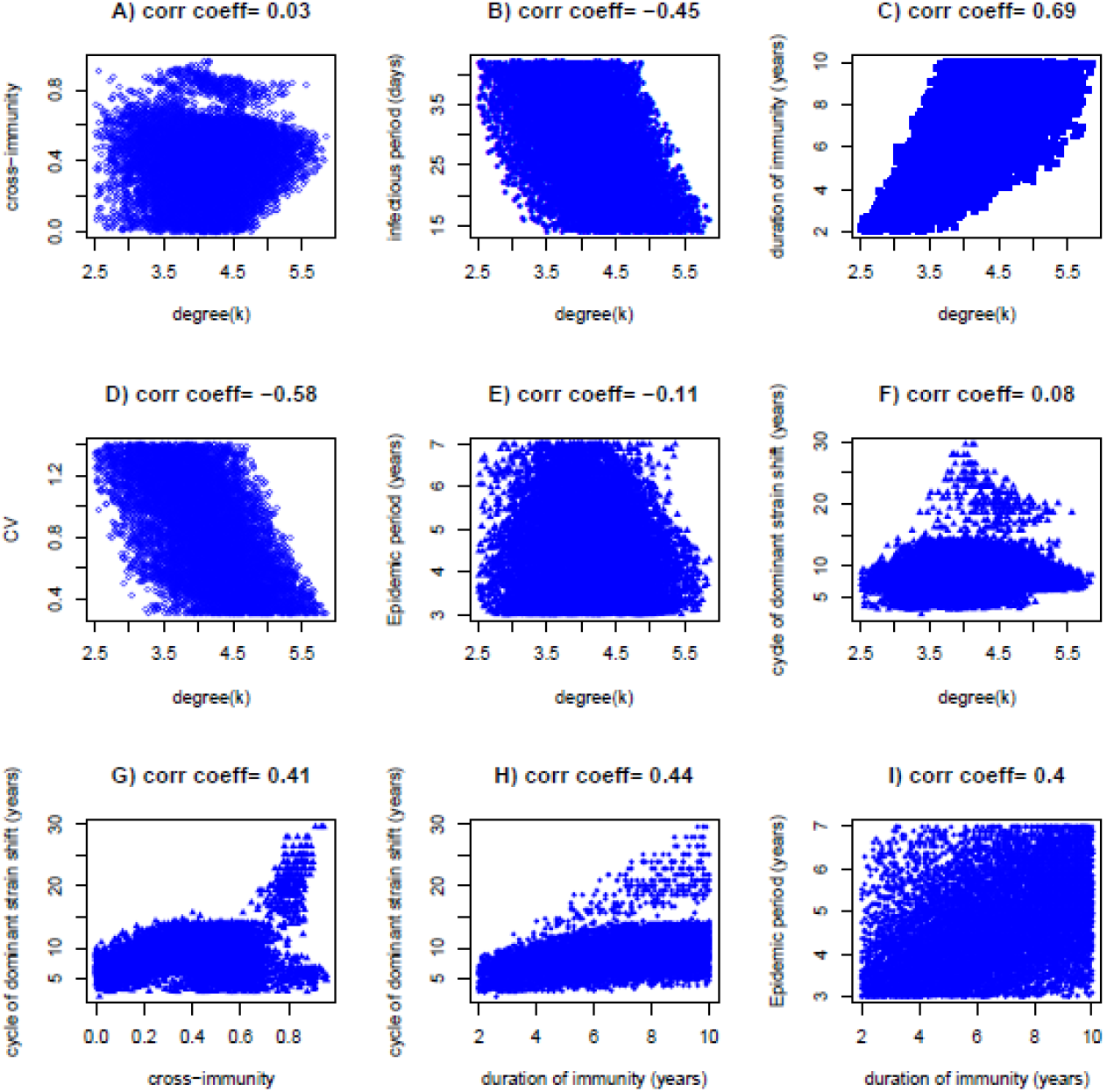
Features of LH sampling of model parameters and dynamic patterns of incidence caused by two asynchronous strains under condition of no interactions within the secondary infection (i.e., *ν* =*μ* =1). An average life span of 70 years and basic reproduction number *R*_0_=1.7 are assumed. As *κ* increases (i.e., network becomes weak), to reproduce epidemic cycles consistent with those observed, the infectious period needs to decrease while the duration of immunity needs to increase.

A value for the interaction parameters *ν* and *μ* of 1.0 implies that there is no interaction on the 2^nd^ infection from the primary infection (This is a special situation considered in Zhang (2016)). If they are less than 1.0, it means the interactions diminish the relevant process. On the contrary, if they are larger than 1.0, they enhance the processes. The ranges listed in Table 1 allow both increasing and decreasing effects to be selected. To constrain the interactions within biologically reasonable limits, we allow both *ν* and *μ* to vary from 0.5 to 2.0. To consider the effect of network-mediated spatial correlation, the contact degree (*κ*) is allowed to vary from 2.5 to 25.

The Runge-Kutta fourth order method was used to solve the model equations (3–4). As our dynamic system is deterministic, there is one dynamical time series under each set of parameter values. For each time series in which infection persists, weekly rates of new infections with each strain are recorded: the first 800 years were discarded and 200 years were used for analysis. To monitor the time series data and calculate the inter-epidemic periods if periodic changes in the incidence of both strains and total number of infections occur, the spectrum function in the R computing language (R Development Core Team. 2015) is employed. The inter-epidemic period (or the duration of epidemic cycle) will be denoted by EpiT. Following Omori *et al*. (2015), the coefficient of variance (CV) of the incidence time series was used to define the shape of epidemic curve and the strength of oscillation in infections over time. Kenri *et al*. (2008) showed that the coefficient of variance (CV) in Japan MP epidemics 1982 to 1990 is about 0.7. In view of this, we regard the epidemics that possess the following characteristics as reasonable approximates to what has been empirically observed in MP epidemics: 0.3≤ CV ≤1.4 and 3 ≤ EpiT ≤ 7 years. In the following we refer to this as the “characteristically recurrent epidemics of MP”.

We study two specific scenarios in relation to the occurrence of MP epidemic cycles. First, assuming that two strains of MP interact only through the cross-immunity during the reinfection process (i.e., *ν*=*μ*=1) we sought to explore how network-structured contacts alone can help build up the characteristics of MP epidemics. Secondly we assume that the primary infection can influence the infectivity and duration of infectiousness of a secondary infection, in addition to the cross-immunity. Under this situation, we examine how these strain interactions can help generate sustained recurrent epidemics and thus relax the requirement of network contacts for the build-up of MP epidemics.

## Results

### Special situation I: Network contact and cross-immunity alone

150,000 combinations of model parameters were sampled with *ν*=*μ* =1 and κranging from 2.5 to 7. Only 11222 combinations generate characteristically recurrent epidemics of MP that are of asynchronous strains and their features are shown in Figure 2. Other 214 combinations generate MP recurrent epidemics that are of synchronous strains (see Appendix A). The results shown in Figure 2A illustrate that reproducing the characteristically recurrent epidemics of MP is not possible unless the contact degree (*κ*) is less than 6.0. That is, without strain interaction within the secondary infection (i.e., *ν*= *μ*= 1), it requires strong network-mediated spatial correlation (c.f., Rozhnova and Nunes. 2009; Zhang 2016) to enable MP epidemic cycling. When cross-immunity is not extremely strong (shown in Figure 2A), two strains asynchronously shift among epidemics; otherwise, they completely synchronise (see Figure A1A in Appendix; c.f., Zhang 2016). The infectious period is negatively correlated while the duration of immunity is positively correlated with the degree of contact (Figure 2B and 2C). This suggests that, all other parameters being equal: within a population of a relatively large contact degree, the infectious period will need to become shorter while the duration of immunity needs to become longer for recurrent epidemics consistent with those observed to occur. Recurrent epidemics generated by a population of small contact degree (*κ* being just larger than 2.5) have a high coefficient of variation and show strong oscillations while those generated by the population of large contact degree (κbeing just less than 6.0) have low coefficient of variation (Figure 2D). Compared to other parameter combinations, both situations result in slightly shorter durations of recurrent epidemics and cycles of dominant strain shift (Figure 2E and 2F), however. Cycles of dominant strain shift are positively associated with the presence of cross-immunity, the duration of immunity and the infectious period (Figure 2G, 2H and 2I), whilst durations of recurrent epidemics are insensitive to these parameters (data not shown).

Two examples of the predicted recurrent epidemics are demonstrated in Figure 3: one is a regular recurrent epidemic and the other irregular. To illustrate the possible mechanisms of oscillation in incidence and the shift of the dominant serotype, we also plot the changes in the susceptible individuals, and the individuals that are immune to strain 1 alone, and to strain 2 alone, and to both strains together. In Figure 3A MP epidemics occur regularly with epidemic period of exactly 4 years and the CV is 0.51. Two strains alternate the dominancy symmetrically from one epidemic to another: when one strain is dominant the other strain remains at extremely low activities. That is, each separate epidemic is mainly caused by one strain. In Figure 3B the duration of recurrent epidemics ranges from 3 years to 5 years with an average of four years. The average CV is 1.08, indicating a strong oscillation comparing to example shown in Figure 3A. The epidemics also vary in the total number of infections. During each epidemic, infections can be due to either mainly one strain or two strains simultaneously.

Comparisons of the upper and bottom graphs in each panel of Figure 3 show that infections oscillate following the changes in proportion of the population that is susceptible. The shift of the dominant strain during the oscillating epidemics is due to changes in the proportion of the population that is immune to different strains: The incidence of one strain will increase when the proportion of people immune to it is low; at the same time the incidence of the other strain will decrease because of the relatively high proportion of the population that is immune. This observation seems to support the hypothesis that a decline in immunity or an increase of the immunologically naïve population may result in the 4-year cycle of epidemic periods (Chalker *et al*., 2011).

### Special situation II: Network contact and strain interactions via cross-immunity during re-infection and interactions within secondary infection

Preliminary sampling experiments indicate that the number of parameter combinations that can generate MP recurrent epidemics decreases quickly as the degree of contacts increases. When the degree of contacts (*κ*) exceeds 15, the combinations of model parameters for MP recurrent epidemics become extremely rare. To save the computational time, the model parameter values were sampled by dividing them into groups by the ranges of *κ* 2.5–6, 6–8, 8–10, 10–12, 12–14, 14–16, 16–17 with respective sampling sizes 200,000, 200,000, 250,000, 250,000, 250,000, 250,000, 500,000. We obtained 23534 combinations of model parameter values that generate characteristically recurrent epidemics of MP that are of asynchronous strains; the maximum of *κ* is 16.2 (see Figure 4A). Other 4426 combinations generate MP recurrent epidemics that are of synchronous strains (see Appendix).

**Figure 4.**
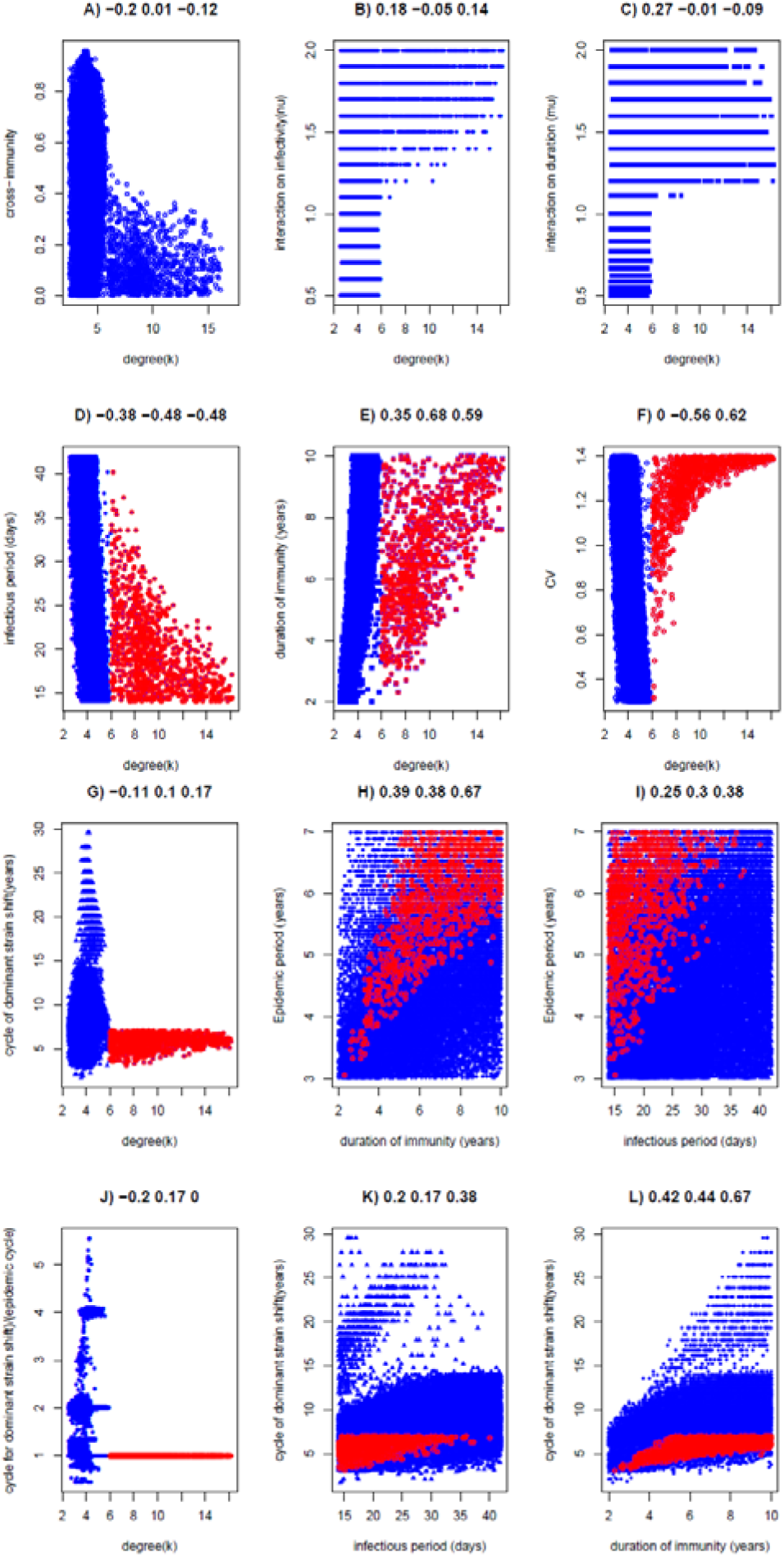
Features of LH samplings of model parameters and dynamic patterns of incidence caused by two asynchronous strains with interactions within the secondary infection. An average life span of 70 years and basic reproduction number *R*_0_=1.7 are assumed. Panel A shows the maximum degree of contacts is 16.2 while panel B and C show that the critical degree for the asynchronous strains is *κ*_ac_= 6.0. In panels D-P) the blue points represent the parameter values when contact degrees *κ*≤*κ*_ac_ and the red points those with contact degrees *κ*>*κ*_ac_. The three values above each panel represent the correlation coefficients between the two variables for all values, the values when *κ*≤*κ*_ac_, and the values when *κ*>*κ*_ac_.

The results shown in Figure 4 indicate that compared to the above situation (I), allowing for strain interactions within secondary infections can lead to the characteristically recurrent epidemics of MP even in a population that has little network mediated spatial correlation. The maximum degree of contacts (16.2) is much larger than the maximum value of 5.9 that was required for situation I. The distributed patterns in Figure 4A, 4B and 4C suggest that there is a critical threshold in the degree of contacts (denoted by *κ*_ac_ for asynchronous strain recurrent epidemics thereafter) separating the mechanisms by which recurrent epidemics consistent with those seen for MP occur. For the parameter values given in Figure 4, *κ*_ac_=6.0. For populations that are of contact degree *κ*<*κ*_ac_, recurrent epidemics occur because of the spatial correlation induced by strong network structure whilst for the populations of relatively loose network structure (*κ*>*κ*_ac_), they occur because of the combination of spatial correlation and strain interactions. We refer them as mechanism 1 and mechanism 2 respectively. It is clearly shown in Figure 4A that although cross-immunity can be any level from 0 to 1 under mechanism 1, only weak cross-immunity levels (<0.4) are required under mechanism 2.

As shown above (special situation I), when *κ* <*κ*_ac_ (mechanism 1), characteristically MP recurrent epidemics are readily generated irrespective of whether the primary infection affects the secondary infection. When strain interactions are present, complicated epidemics can be generated (see Figure 5). Under the loose network structure (*κ*>*κ*_ac_) (mechanism 2), MP recurrent epidemics can be produced only when the primary infection enhances the infectivity (*ν*>1) (Figure 4B) and prolongs the infectious period (*μ* >1) (Figure 4C) of the secondary infection. Conversely, if strain interactions diminish the transmissibility of a secondary infection, shifts in the dominant strain in epidemics cannot occur. As in special situation I, relatively short infectious periods of MP at populations of high contact degree are required while relatively longer durations of immunity are needed to generate the recurrent MP epidemics (Figure 4D and 4E). The conditions for the emergence of synchronous strains are different and are shown in Appendix.

**Figure 5.**
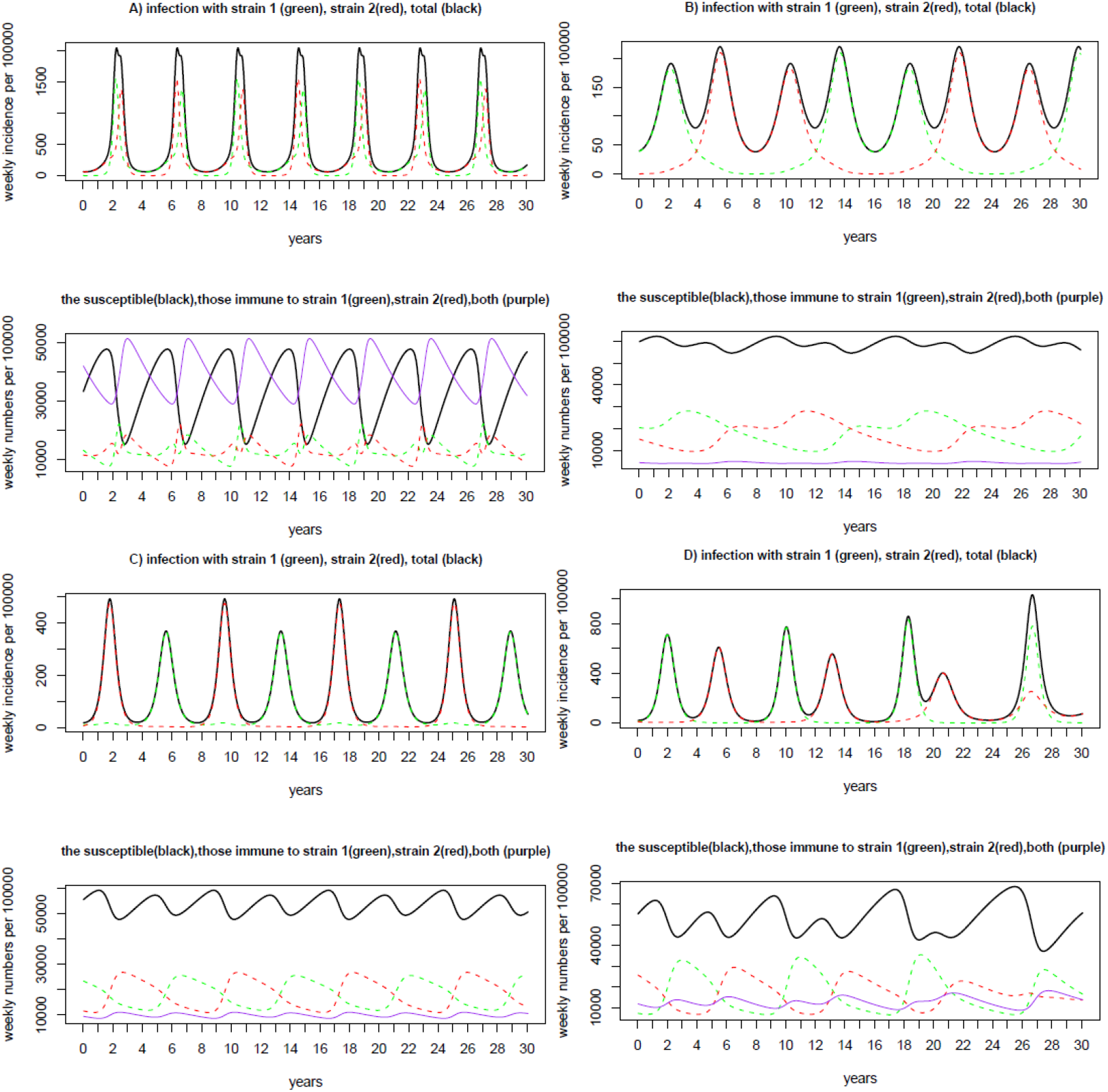
Four examples of epidemic curve from LHS samples that generate MP recurrent epidemics with strain interactions within the secondary infection. Panel A) *κ*=8.79, *ψ* =0.012, *D*=15.4 days, *d* =4.3 years, *ν*=1.8, *μ*=1.43, and EpiT =4 years, CV =1.16; B) *κ* =5.04, *ψ*=0.7, *D* =14.4 days, *d*=8.6 years, *ν*=1.4, *μ*=0.59, and EpiT =4 years, CV =0.479; C) *κ*=5.02, *ψ*=0.424, *D*=14.1 days, *d* =8.8 years, *ν*=1.2, *μ*=0.77, and EpiT=4 years, CV =0.896; *D*) *κ*=3.67, *ψ*=0.185, *D* =32.6 days, *d* =5.4 years, *ν*=1.6, *μ*=0.63, and EpiT =4 years, CV =0.598. The legend is provided in the title to each figure.

Dynamic patterns including the shape of oscillations and durations of recurrent epidemics and cycle of dominant strain shift are shown in Figure 4F-4L. Figure 4F shows that the shape of the epidemic curve (i.e., the coefficient of variance (CV) of incidence along the time) is positively correlated with *κ* when *κ* >*κ*_ac_, while CV decreases with *κ* when *κ*≤*κ*_ac_. This suggests that within a looser networked population, the oscillation in incidence tends to become stronger. However, CV is not sensitive to the other model parameters (data not shown).

The cycle of dominant strain shift ranges from 3 to 30 years (Figure 4G), which covers the observational ranges: 10–16 years (Kendri *et al*., 2008; Kogoj *et al*., 2017). Figure 4J indicates that when *κ*<*κ*_ac_, the cycle of dominant strain shift can be 1–6 times the duration of recurrent epidemics; when *κ* >*κ*_ac_, the cycle of dominant strain shift approximates the epidemic cycle. Both the duration of recurrent epidemics and cycle of dominant strain shift are positively associated with the duration of immunity, especially under mechanism 2 (Figure 4H and 4L). Under mechanism 2, they are weakly and positively associated with the infectious period (Figure 4I and 4K). Otherwise, they are insensitive to other parameters (see Figure 4G for the relationship between cycle of dominant strain shift and degree of contacts).

Four typically recurrent epidemic examples are illustrated in Figure 5. They show different oscillation patterns. In panel A) two strains are of comparable activity levels with the strain that starts early dominating the epidemics; strain dominancy alternates regularly among epidemics cycle and oscillate with the same period of four years. In panel B) two strains shift dominancy with each strain dominating two epidemics consecutively before switching strain dominancy; the two consecutive epidemics are mainly activated by the dominant strain while the other strain remains at very low activity. In panel C) although epidemics take place regularly, recurrent epidemics consist of two different epidemics: one with high peak and narrow active period, the other with lower peak but wide active period; two strains alternate their dominancy accordingly. In panel D) two strains alternate with irregular peaks and total incidence within each epidemic. The diverse patterns may mimic real observations in MP epidemics (Kenri *et al*., 2008; Brown *et al*., 2016).

Comparing the levels of infection and of immunity can shed light on the underlying mechanisms of recurrent epidemics. It is obvious from the Figure 5 that the proportion of individuals that are simultaneously immune to both strains is kept low except for panel A) where it oscillates within a wide range and anti-correlates with the proportion susceptible. The proportion of individuals immune to one strain is temporally highly anti-correlated with the proportion of these immune to the other strain, with absolute correlation coefficients >80% for panels B), C) and D); while for panel A) they are weakly correlated. This difference reflects their different levels of cross-immunity. Panels A) and B) show predictions obtained for a situation in which the primary infection strain increases the infectivity of secondary strain (*ν*>1). In panel A) although the dominant strain shifts between among epidemics, the difference in the proportion immune or susceptible to the dominant and non-dominant strains is small. In panel B), one strain is dominant while the other remains at a very low incidence, which continues over further epidemics even if the proportion of individuals that are immune to the strain exceeds the proportion that is immune to the other strain. The dominancy only changes when the difference in immunity to a given strain increases substantially. So under this situation, every strain dominates continuously over two epidemics before the strain dominancy switches. As found for situation I, oscillations in the infection incidence follow changes in the proportion of the population that is immune and susceptible. The change of dominant strain during the recurrent epidemics is due to the exchange in immunity to different strain: the increase in infection activity of one strain follows the relatively low immunity to the strain (Chalker *et al*., 2011).

The findings assuming values for *R*_0_ of 1.3 and 1.5 are similar to those obtained assuming a value of 1.7, although the critical threshold value in the degree of contacts differs (see Figure 6). Under low values for *R*_0_ (1.3), for example, the critical degree of contact decreases to *κ*_ac_=4.7 and the maximum contact degree decreases to 7.4.

**Figure 6.**
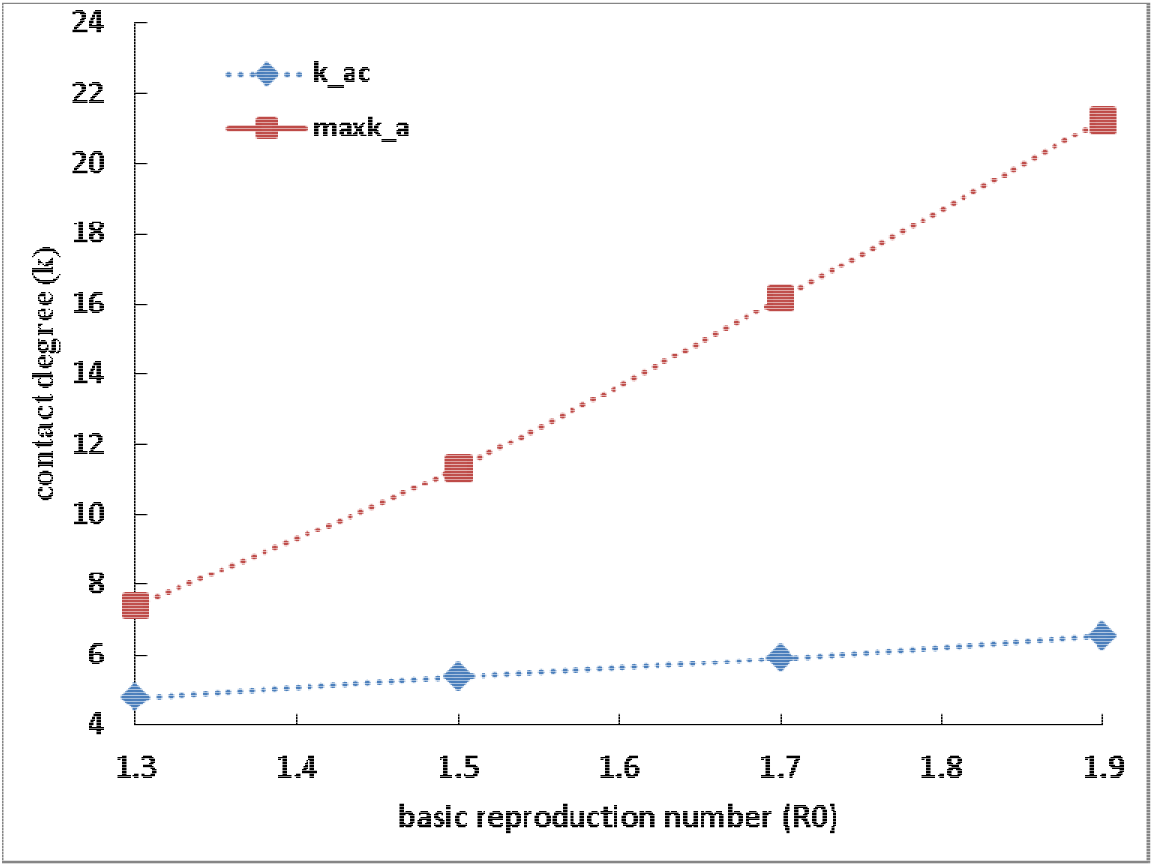
Critical threshold degrees of contacts under different transmissibility

## Discussion

In this study we demonstrate that spatial correlation mediated by contact network of human population and positive strain interactions within secondary infection work cooperatively to drive MP infection incidence into recurrent epidemics occurring every three to seven years with dominant strains shifting among epidemics.

Accounting for realistic host population structure in infectious disease modelling is important. It has been recognised that it is necessary to take true network contacts among human populations to explain the observed dissemination patterns of infectious disease (e.g., Brockmann and Helbing, 2013). The results shown in Figure 2 where no strain interaction is assumed illustrate that contact networks of degree less than 6 are required for the recurrent epidemics that occur every 3–7 years with alternation of dominant strain. A key property of a network is its degree distribution. For community, school and hospital networks, empirical studies suggest that the average degree is 6.5 (Leventhal *et al*., 2015). This empirical information of human contact networks therefore suggests that the network model without strain interaction within secondary infection could not be a candidate mechanism for MP recurrent epidemics.

Interaction between different strains during re-infection (i.e. cross-immunity) is well known and has attracted much effort to study and measure it. Once an individual is re-infected by another strain, are there any interactions between the primary infection strain and the secondary infection strain within the secondary infection? Surely recovery from primary infection will not leave immunocompetent host individuals naïve. It is theoretically reasonable to argue that the non-naïve individuals would have other changes which might in some ways alter the secondary infection by other strains (see Zhang and Cao, 2014 for more general reasoning). As MP parasitizes the respiratory tract epithelium of humans, the primary infection with one strain can, for example, damage the airway (Song *et al*., 2015), which could then alter the ecological niche of the secondary infection strain. Further, the primary infections of the upper or lower respiratory tract can be followed by extrapulmonary complications (Tsiodras *et al*., 2005). As far as the transmission dynamics are concerned, the modifications in the non-naïve individuals might change the infectivity and duration of secondary infection. To our knowledge, we have not found any clear empirical data for these interactions among strains of MP, although this may reflect a lack of research into MP pathogenicity. In principle, prior exposure of an individual to a strain could have no effect or either decrease or increase the individual’s ability to clear an infection with a differing strain, with potential to increase or reduce the overall transmission. The theoretical analysis in this study shows that only positive strain interactions increase the infectiousness of a secondary infection to facilitate the generation of recurrent MP epidemics. It is expected that the experimental observations and measurement of strain interactions within secondary infections will provide vital proof to support or disprove the combination of network mediated spatial correlations and strain interactions with secondary infections as a determinant of epidemic recycling of MP.

We found that there is a positive association between the durations of recurrent epidemics and the duration of immunity (Figure 4H). This finding is consistent with that of Goncalves *et al*. (2011) but differ from that of Omori *et al*. (2015). Further, the cycle of dominant strain shift also shows positive correlations with the duration of immunity (Figure 4L). These positive correlations become stronger under the mechanism whereby recurrent epidemics are generated by the combination of network mediated spatial correlations and strain interactions with secondary infection. Under this mechanism, both cycles are intermediately linked with the infectious period (Figure 4I and 4K). In contrast to the observations of Keeling and Rohani (2008) and Omori *et al*. (2015), the cycle of the dominant strain shift is insensitive to cross-immunity.

Despite our simplifying assumptions, the network model of the transmission dynamics of two strains presented here remains complicated. The distributions of both infectious period and the duration of immunity are implicitly assumed to be exponential. Omori *et al*. (2015) suggested that assuming that the duration of immunity follows a distribution with a variance about 0.63, which is much smaller than that of the exponential distribution, models that assume that people mix randomly can produce the periodicity of MP recurrent epidemics. Can the reduced variation in the distribution of the duration of immunity help build up the recurrent epidemics in our network model? To see this, we construct a SI1I2R1R1R2R2J1J2RR network model by separating recovery stages into two equal parts. Therein the immunity period follows a gamma distribution of shape parameter =2. This model has 11 nodes and is described by 65 differential equations. Nonetheless, the simulations (data not shown) show that this more complicated model does not give any noticeably different results. It is a technical challenge in our network model to generate gamma distributed duration of immunity that is comparable to that required in Omori *et al*. (2015).

We modelled human population structure as a static, unweighted network wherein each individual has an equal number of links with other people. The real-world contacts between individuals are dynamic and the network degree of contacts varies from person to person (e.g., Guclu *et al*., 2016). How these heterogeneities in contact networks affect the model results, albeit being worth further analysing, is an analytically and computationally challenging issue.

In this study we constrained both the interaction parameters v and μ, which describe the effects of primary infection strain on the infectivity and duration of infection by secondary strain respectively, at the ranges from 0.5 to 2.0. If we had widened their ranges, requirement for the limited contact degree is further reduced. This is in agreement with the previous studies (Zhang and Cao, 2014): under strong strain interactions alone, epidemic cycling becomes possible even under assumptions of the homogenous mixing.

In conclusion, we have illustrated that multiple strains that co-circulate within a network structured population and interact positively as secondary infections with primary infections generate the MP epidemics of 3–7 year interval and alternating dominant strains. This model supports the theory that epidemic shifts in MP may be attributed to population immunity not only to the immunogenic strain in question, but also with the influence of cross protection and other enhanced effects from the second strain type and that transmission via patient networks within the population combine to produce MP epidemic cycles. Though the strain interactions within a secondary infection are theoretically possible, currently no reliable evidence exists to suggest whether either a positive or negative strain interaction occurs. We hope this study can encourage experimental studies to detect and measure interactions between strains of MP. This will benefit our understanding of MP and provide crucial information for us to predict and thus control its recurrence.

## Acknowledgments

The work was carried out whilst the authors were employed by Public Health England. We thank Elaine Stanford for her kind support for this work.

## References

Anderson RM and May RM. 1991. Infectious disease of humans: dynamics and control. Oxford: Oxford University Press.

Brockmann D, Helbing D. 2013. The hidden geometry of complex, network-driven contagion phenomena. Science 342:1337–1342. doi: 10.1126/science.1245200

Brown RJ, Holden MT, Spiller OB, Chalker, VJ. 2015. Development of a multilocus sequence typing scheme for molecular typing of *Mycoplasma pneumoniae*. Journal of Clinical Microbiology 53:3195–3203. doi:10.1128/JCM.01301-15

Brown RJ, Nguipdop-Djomo P, Zhao H, Stanford E, Brad Spiller O, Chalker VJ. 2016. Mycoplasma pneumoniae Epidemiology in England and Wales: A national perspective. Frontiers in Microbiology 7:157. doi:10.3389/fmicb.2016.00157

Chalker VJ, Stocki T, Mentasti M, Fleming D, Harrison TG. 2011. Increased incidence of Mycoplasma pneumoniae infection in England and Wales in 2010: multiocus variable number tandem repeat analysis typing and macrolide susceptibility. Euro Surveillance 16(19):pii=19865. Available online: http://www.eurosurveillance.org/ViewArticle.aspx?ArticleId=19865

Clyde WA. 1993. Clinical overview of typical Mycoplasma-Pneumoniae infections. Clinical Infectious Diseases 17: S32–S36.

Diekmann O, Heesterbeek H, Britton T. 2013. Mathematical tools for understanding infectious disease dynamics. Princeton University Press, Princeton and Oxford.

Dorigo-Zetsma JW, Dankert J, Zaat SA. 2000. Genotyping of Mycoplasma pneumoniae clinical isolates reveals eight P1 subtypes within two genomic groups. Journal of Clinical Microbiology 38: 965–970. PMID: 10698981

Dumke R, von Baum H, Lück PC, Jacobs E. 2010. Subtypes and variants of Mycoplasma pneumoniae: local and temporal changes in Germany 2003-2006 and absence of a correlation between the genotype in the respiratory tract and the occurrence of genotype-specific antibodies in the sera of infected patients. Epidemiology and Infection 138(12):1829–1837. https://doi.org/10.1017/S0950268810000622

Dumke R, Jacobs E. 2011. Culture-independent multi-locus variable-number tandem-repeat analysis (MLVA) of Mycoplasma pneumoniae. Journal of Microbiological Methods 86(3):393–396. doi:10.1016/j.mimet2011.06.008

Dumke R, Strubel A, Cyncynatus, C, Nuyttens H, Herrmann R, Lück C, Jacobs E. 2012. Optimized serodiagnosis of *Mycoplasma pneumoniae* infections. Diagnostic Microbiology and Infectious Disease 73:200–203. doi:10.1016/ j.diagmicrobio.2012. 02.014

Dumke R, Jacobs E. 2016. Antibody Response to *Mycoplasma pneumoniae*: Protection of Host and Influence on Outbreaks? Frontiers in Microbiology 7:39. doi:10.3389/fmicb. 2016.00039

Eames KTD, Keeling MJ. 2002. Modelling dynamic and network heterogeneities in the spread of sexually transmitted diseases. Proc Natl Acad Sci USA 99:13330–13335. https://doi.org/10.1073/pnas.202244299

Foy HM, Kenny GE, Sefi R, Ochs HD, Allen ID. 1977. Second attacks of pneumonia due to Mycoplasma pneumoniae. Journal of Infectious Disease 135:673–677. PMID:856921

Foy HM. 1993. Infections caused by *Mycoplasma pneumoniae* and possible carrier state in different populations of patients. Clinical Infectious Diseases 17:S37–S46. doi:10.1093/clinids/17.Supplement_1.S37

Goncalves S, Abramson G, Gomes MFC. 2011. Oscillations in SIRS model with distributed delays. European Physical Journal B 81:363–371. https://doi.org/10.1140/epjb/e2011-20054-9

Guclu H, Read J, Vukotich Jr CJ, Galloway DD, Gao H, Rainey JJ, Uzicanin A, Zimmer SM, Cummings DA. 2016. Social Contact Networks and Mixing among Students in K-12 Schools in Pittsburgh, PA. PLOS One 11(3):e0151139. doi:10.1371/journal.pone. 0151139

Iman RL, Helton JC, Campbell JE. 1981. An approach to sensitivity analysis of computer models, Part 1. Introduction, input variable selection and preliminary variable assessment. Journal of Quality Technology 13(3):174–183.

Ito I, Ishida T, Osawa M, Arita M, Hashimoto T, Hongo T, Mishma M. 2001. Culturally verified *Mycoplasma pneumoniae* pneumonia in Japan: a long-term observation from 1979–99. Epidemiology and Infection 127:365–367. doi: 10.1017/s0950268801005982

Jacobs E. 2012. *Mycoplasma pneumoniae*: now in the focus of clinicians and epidemiologists. Euro Surveillance 17:20084. http://www.eurosurveillance.org/ViewArticle.aspx?ArticleId=20084

Kenri T, Okazaki N, Yamazaki T, Narita M, Izumikawa K, Matsuoka M, Suzuki S, Horino A, Sasaki T. 2008 Genotyping analysis of Mycoplasma pneumoniae clinical strains in Japan between 1995 and 2005: type shift phenomenon of M. pneumoniae clinical strains. Journal of Medical Microbiology 57:469–475. doi:10.1099/jmm.0.47634-0

Keeling MJ, Rohani P. 2008. Modelling Infectious Disease in Humans and Animals. Princeton. NJ, USA: Princeton University Press.

Kogoj R, Praprotnik M, Mrvič T, Korva M, Kese D. 2017. Genetic diversity and macrolide resistance of Mycoplasma pneumoniae isolates from two consecutive epidemics in Slovenia. European Journal of Clinical Microbiology and Infectious Diseases 37:99–107. https://doi.org/10.1007/s10096-017-3106-5

Korppi M, Heiskanen-Kosma T, Kleemola M. 2004. Incidence of community-acquired pneumonia in children caused by *Mycoplasma pneumoniae*: serological results of aprospective, population-based study in primary healthcare. Respirology 9:109–114.doi:10.1111/j.1440- 1843.2003.00522.x

Leventhal GE, Hill AL, Nowak MA, Bonhoeffer S. 2015. Evolution and emergence of infectious diseases in theoretical and real-world network. Nature Communications 6:6101. doi:10.1038/ncomms7101 (2015).

Lind K, Benzon MW, Skov Jensen J, Clyde WA. 1997. A seroepidemiological study of *Mycoplasma pneumoniae* infections in Demark over 50-year period 1946-1995. European Journal of Epidemiology 13:581–586. https://doi.org/10.1023/A:1007353121693

Martinez MA, Ruiz M, Zunino E, Luchsinger V, Aguirre R, Avendano LF. 2010. Identification of P1 types and variants of Mycoplasma pneumoniae during an epidemics in Chile. Journal of Medical Microbiology 59:925–929. doi: 10.1099/jmm.0.018333-0.

Morozumi M, Iwata S, Hasegawa K, Chiba N, Takayanagi R, Matsubara K, Nakayama E, Sunakawa K, Ubukata K. 2008. Increased macrolide resistance of *Mycoplasma pneumoniae* in paediatric patients with community acquired pneumonia. Antimicrob Agents Chemother 52:348–350. doi: 10.1128/AAC.00779-07

Mossong J, Hens N, Jit M, Beutels P, Auranen K, Mikolajczyk R, Massari M, Salmaso S, Tomba GS, Wallinga J, Heijne J, Sadkowska-Todys M, Rosinska M, Edmunds WJ. 2008. Social contacts and mixing patterns relevant to the spread of infectious diseases. PLoS Medicine 5:e74. https://doi.org/10.1371/journal.pmed.0050074

Nakata Y, Omori R. 2015. Delay equation formulation for an epidemic model with waning immunity: an application to mycoplasma pneumoniae. IFAC-PapersOnLine 48:132–135. https://doi.org/10.1016/j.ifacol.2015.11.024

Nguipdop-Djomo P, Fine PEM, Halsby KD, Chalker VJ, Vynnycky E. 2013. Cyclic epidemics of *Mycoplasma pneumoniae* infections in England and Wales from 1975 to 2009: time-series analysis and mathematical modelling. Lancet 382:S78. doi:10.1016/S0140-6736(13)62503-9

Omori R, Nakata Y, Tessmer HL, Suzuki S, Shibayama K. 2015. The determinant of periodicity in Mycoplasma pneumoniae incidence: an insight from mathematical modelling. Scientific Reports 5:14473. doi:10.1038/SREP14473

Pastor-Satorras R, Vespignani A. 2001. Epidemic spreading in scale-free networks. Physical Review Letter 86:3200–3. doi:10.1103/PhysRevLett.86.3200

Pereyre S, Charron A, Hidalgo-Grass C, Touati A, Moses AE, Nir-Paz M, Bebear C. 2012. The Spread of Mycoplasma pneumoniae Is Polyclonal in Both an Endemic Setting in France and in an Epidemic Setting in Israel. PLoS ONE 7(6): e38585. doi:10.1371/journal.pone.0038585

R Development Core Team. 2015. R: a language and environment for statistical computing. https://www.r-project.org/

Rozhnova G, Nunes A. 2009. Fluctuation and oscillations in a simple epidemic model. Physical Review E 79:041922. doi:https://doi.org/10.1103/PhysRevE.79.041922

Simmons WL, Daubenspeck JM, Osborne JD, Balish MF, Waites KB, Dybvig K. 2013. Type 1 and type 2 strains of Mycoplasma pneumoniae form different biofilms. Microbiology 159:737–747. doi:10.1099/mic.0.064782-0.

Song Q, Xu B-P, Shen K-L. 2015. Effects of bacterial and viral co-infections of mycoplasma pneumoniae in children: analysis report from Beijing Children’s hospital between 2010 and 2014. International Journal Clinical and Experimental Medicine 8(9):15666–15674. PMID:26629061

Spuesens EB, Oduber M, Hoogenboezem T, Sluijter M, Hartwig NG, van Rossum AM, Vink C. 2009. Sequence variations in RepMP2/3 and RepMP4 elements reveal intragenomic homologous DNA recombination events in Mycoplasma pneumoniae. Microbiology 155:2182–2196. PMID:19389769

Suzuki Y, Seto J, Itagaki T, Aoki T, Abiko C, Matsuzaki Y. 2015. Gene mutations associated with macrolide-resistance and p1 genotyping of *Mycoplasma pneumoniae* isolated in Yamagata, Japan, between 2004 and 2013. Kansenshogaku Zasshi 89:16–22. https://doi.org/10.11150/kansenshogakuzasshi.89.16

The World Fact-book Life Expectancy. Cia.gov. 2012. https://www.cia.gov/library/publications/the-world-factbook/fields/2102.html

Tsiodras S, Kelesidis I, Kelesidis T, Stamboulis E, Giamarellou H. 2005. Central nervous system manifestations of Mycoplasma pneumoniae infections. Journal of Infection 51(5):343–354. doi:10.1016/j.jinf.2005.07.005

Waites KB, Talkington DF. 2004. Mycoplasma pneumoniae and its role as a Human pathogen. Clinical Microbiology Reviews 17(4):672–728. doi:10.1128/CMR.17.4.697-728.2004

Watts DJ, Strogatz SH. 1998. Collective dynamics of small world networks. Nature 393:440–442. doi:10.1038/30918

Winchell JM. 2013. *Mycoplasma pneumoniae* – A national public health perspective. Current Pediatric Reviews 9(4). doi:10.2174/15733963113099990009

Woodhead M, Macfarlane J. 2000. Local antibiotic guidelines for adult community-acquired pneumonia (CAP): a survey of UK hospital practice in 1999. Journal of Antimicrobial Chemotherapy. 46:141–143. PMID:10882705

Zhang X-S. 2016. Epidemic cycling in a multi-strain SIRS epidemic network model. Theoretical Biology and Medical Modelling 13:14 doi:10.1186/s12976-016-0040-7

Zhang X-S, Cao K-F. 2014. The Impact of Coinfections and Their Simultaneous Transmission on Antigenic Diversity and Epidemic Cycling of Infectious Diseases. BioMed Research International 375862 doi:10.1155/2014/375862

Zhao F, Liu G, Wu J, Cao B, Tao X, He L, Meng F, Zhu L, Lv M, Yin Y, Zhang J. 2013. Surveillance of macrolide-resistant *Mycoplasma pneumoniae* in Beijing, China, from2008 to2012. Antimicrob. Agents Chemother. 57:1521–1523. doi:10.1128/AAC. 02060-12

Zhao F, Liu L, Tao X, He L, Meng F, Zhang J. 2015. Culture-independent detection and genotyping of *Mycoplasma pneumoniae* in clinical specimens from Beijing China. PLoS ONE 10:e0141702.doi: 10.1371/journal.pone.0141702

